# Mosquito Dispersal in Context

**DOI:** 10.1101/2025.03.22.642900

**Authors:** Héctor M. Sánchez C., Sean L. Wu, John M. Henry, Carlos A. Guerra, David S. Galick, Guillermo A. Gárcia, John M. Marshall, David L. Smith

**Affiliations:** Divisions of Biostatistics & Epidemiology, University of California, Berkeley, Berkeley, California, USA; Institute for Health Metrics and Evaluation, University of Washington, Seattle, WA; Merck & Co., Inc., Rahway, NJ, USA; Seattle University, Seattle, Washington; MCD Global Health, Silver Spring, Maryland, USA; Department of Health Metrics Sciences, School of Medicine, University of Washington, Seattle, WA

## Abstract

Mosquito dispersal plays an important role in mosquito ecology and mosquito-borne pathogen transmission. While reaction-diffusion and patch-based models with simple flux assumptions for emigration have played a predominant role in modeling mosquito dispersal, mosquito behavioral ecology – in particular, the process of searching for resources – is usually ignored by diffusion-based models. We thus set out to analyze mosquito movement using highly mimetic models, to see what we could learn from a different approach. Here, we present two behavioral state microsimulation models, in which mosquitoes are in behavioral states and search for required resources, which are distributed on a landscape. Models of this sort are laborious and challenging to work with, so we developed **ramp.micro**, an R package to build, solve, analyze, and visualize behavioral state microsimulation models for mosquitoes. Mosquito population dispersal is an emergent feature in these complex systems, so we developed new methods to describe and understand it. We show that population structure might not be revealed by a crude analysis of observed mosquito movement patterns, unless it can somehow account for sequences of flights for mosquitoes in behavioral states trying to accomplish a goal. We also show that even when resources are distributed randomly and uniformly, mosquito populations tend to form highly spatially structured communities. We also show that some heterogeneity in mosquito population densities is attributed to features of a network defined by searching and the spatial distribution of resources. These models highlight the importance of understanding mosquito behavioral states and resource availability in understanding mosquito population dispersal, and they point to the importance of local context. Motivated by these dynamics, spatial models for mosquito ecology and mosquito-borne pathogen transmission would benefit from considering resource availability as a factor affecting mosquito movement and dispersal.

**Author summary:** We develop highly realistic population dynamic models for mosquito behavioral ecology where mosquitoes move around on landscapes made up of resources located at points in space. The mosquitoes are in behavioral states – blood feeding, egg laying, or sugar feeding – and they end each search at a point where the resource they need can be found. Even with simple assumptions, mosquito populations can be highly structured. Because mosquitoes tend to leave areas that lack a resource, and they tend to stay in areas that have all required resources, the scarcest resources end up determining the structure of mosquito populations.

## Introduction

Hundreds of pathogen species are transmitted among vertebrate hosts by mosquitoes. Mosquito-borne pathogen transmission, the transfer of pathogens through blood feeding resulting in infection, includes two kinds of events: from an infectious mosquito into the blood of a susceptible vertebrate host (through the bite, in saliva); or from an infectious host into the gut of a mosquito (in the blood meal). Pathogen dispersal involves pathogen transmission among infected mosquitoes or vertebrate hosts that move around after getting infected [1]. Mosquito dispersal is thus important for pathogen transmission dynamics as well as mosquito spatial ecology, population dynamics, and population genetics [2, 3]. Mosquito dispersal affects the spread of genetically modified mosquitoes [4], mosquito population dynamic responses to vector control [5, 6], area effects of vector control [7, 8], and the size and clusters and buffer zones for vector control field trials [9]. A core challenge for mosquito ecology and the study of mosquito-borne pathogen dispersal has been quantifying dispersal, often using mark-release-recapture studies [10]. The spatial transmission dynamics of pathogens and mosquito spatial ecology have been studied using a variety of techniques and methods, including mathematical models [10–13]. Most mathematical models of mosquito or pathogen dispersal have been diffusion-based [11], Diffusion is a phenomenological approach that ignores most aspects of mosquito biology, including mosquito searching and flight bouts, the emerging patterns of mosquito population dispersal, and heterogeneous transmission dynamics [2, 14, 15]. An alternative mathematical approach to studying mosquito population movement is through analysis of behavioral state, micro-simulation models, which are highly mimetic; mosquito dispersal is modeled as a sequence of intentive flight bouts ending at a point in space where one of two or more resources can be found [12, 15–18]. To lower the costs of setting up and analyzing such models and to make the study of such models more replicable, we developed an R package hosted on GitHub called ramp.micro. Here, using ramp.micro, we study mosquito dispersal and potential pathogen dispersal by mosquito populations among spatially distributed resources using behavioral state, micro-simulation models. Using methods from network analysis, we show how resource co-distributions give rise to complicated population structure on invented landscapes.

On one level, a mechanistic understanding of mosquito movement could focus on how an individual mosquito searches for, detects, and approaches a single resource. Such studies would link mosquito physiology to the associated behaviors, such as the tactical algorithm that a mosquito uses to locate a resource by following an odor plume upwind [19]. In such studies, the physiological state of a mosquito affects what it senses, and sensory input is linked to a search algorithm that determines how a mosquito reacts to proximal cues designed to move the mosquito towards a resource [20–22]. These are complex behavioral algorithms: mosquitoes respond to environmental cues such as wind speed and relative humidity, and they are attracted to resources at varying distances by odors, visual cues, and CO_2_ [21, 22]. Mosquitoes must navigate around barriers and through a matrix of heterogeneous habitats [2]; and finally, they must use the resource.

At another level, mosquitoes need several resources, and each resource could be available at multiple different locations [2, 3, 17]. The study of mosquito dispersal is thus not just about how mosquitoes fly to a resource, but how they end up at one location among many that are possible. Aggregate movement patterns of mosquito populations arise from behavioral algorithms responding to the spatial distribution of resources on a landscape in a sequence of flight bouts, in which the destination of one search bout is the beginning of another [15, 17, 23, 24]. Some required resources are obvious. New mosquitoes must find a mate, and they may need vertebrate blood or sugar before they start producing eggs. After mating and taking a blood meal, a mosquito must find aquatic habitats to lay eggs. Thereafter, the adult female mosquito’s behavior cycles through blood feeding, egg laying, sugar feeding, and perhaps other behavioral modalities to find and use other resources. The endpoint of one successful search becomes the starting point for another, as a mosquito’s physiological state changes after consuming or using a resource and its behavior reorients towards finding another. Throughout the search process, mosquitoes must rest, so resting habitats are essential way stations. The co-distribution of resting sites and resources forms a template that structures movement. Dispersal of mosquito populations depends on context [12, 15, 23].

Among mathematical models describing mosquito population movement, the predominant formalism has been diffusion. While simple diffusion-based models have been widely used, mosquito flight behavior – as we have just described it – violates the basic assumptions of diffusion models in several ways, particularly searching for resources through a cycle of blood feeding and egg laying. Mosquito flight bouts are generally considered to be either migratory, appetential or consumatory [2, 3]. Mosquito appetential and consumatory flights have a purpose – to find and consume resources – so mosquito movement involves a sequence of intentive flights [3, 25]. Since mosquito behavioral states change, a mosquito could seek a different resource during each flight bout. While mosquito flight paths during search may be modeled as a kind of rough diffusion [13], such models are not designed to understand aggregate movement patterns of mosquito populations in terms of the endpoints of sequences of flights moving among two or more resources [15, 17, 23]. Mosquitoes can use wind to search, and there are good reasons to believe that mosquitoes are opportunistic – they will tend to use the resources that are closer to where they start searching, but flight distances are not always short. With the help of wind, a single mosquito could move a very long distance in a single flight bout [3, 15, 26]; a sequence of mosquito flights could resemble a Levy walk [27]. Alternative mathematical approaches may be more appropriate for understanding how the aggregate movement patterns of mosquito populations could be affected by the distribution of resources, mosquito search algorithms, wind, and other factors. Dispersal of mosquito populations might be understood, not as diffusion, but as random walks on directed graphs (search and dispersal) among nodes that represent the spatial distributions of resources.

Here, we analyze dispersal using a mathematical formalism involving graphs, derived from simulating population dynamics through flight bouts to search for and use resources represented as point sets [12, 15, 17, 18]. The mosquitoes are in behavioral states, and we simulate behavioral state transitions of mosquitoes as they search for and move among two or more heterogeneously distributed resources [15, 23]. We couple adult and immature mosquito populations: adults move around to blood feed, sugar feed, and lay eggs [1, 15]; then mosquito eggs hatch and immature mosquito populations develop in aquatic habitats until they emerge as adults. To simulate search and flight, we formulate rules that determine how mosquitoes select a nearby resource. We analyze mosquito movement in various ways, including the dispersal of eggs away from a natal habitat, and potential dispersal of pathogens away from a point of infection – a metric that is closely related to vectorial capacity [15]. Dispersal patterns among points can be represented as weighted graphs, so we have used concepts from network analysis to analyze the resulting patterns. We extend this analysis to understand how movement patterns would affect mosquito ecology and pathogen transmission.

## Results

We developed two classes of discrete-time, adult behavioral state, micro-simulation models; and we also developed discrete-time models for immature mosquito populations. The two components are linked through egg laying by adults and emergence of adults from aquatic habitats. Mathematical details of the models are described in Methods, and the notation is summarized in Table 1.

**Table 1.**
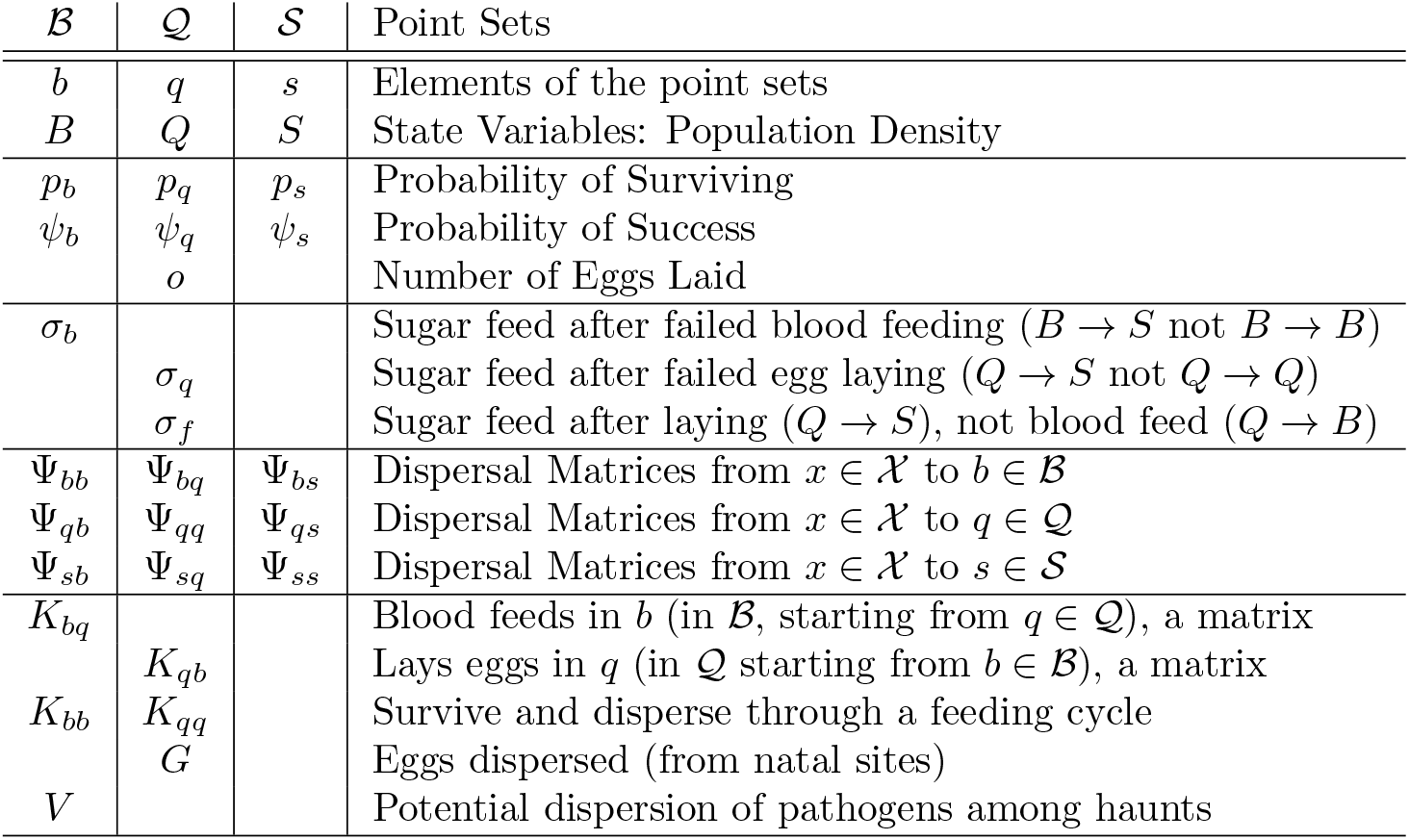
The notation, including variables, parameters, and terms for adult population models and terms describing dispersal. The first column is blood feeding; the second egg laying; and the third sugar feeding.

In adult mosquito models, the adult mosquito population is sub-divided into a set of mutually exclusive and collectively exhaustive compartments where each compartment represents a different physiological or behavioral state. The locations of resources are point sets: aquatic habitats for egg laying, denoted 𝒬; haunts for blood feeding, ℬ; sugar sources for sugar feeding, 𝒮. We call these point sets the *resource landscape*.

Each day, some fraction of the mosquitoes makes a search attempt to find resources. The outcome depends on their state and location. At the end of that search attempt bout, a mosquito has either died, failed, or succeeded. State and location specific parameters describe the fraction surviving each time step (*p*), and the fraction that succeeds in finding and using a resource (*ψ*). In some cases, we assume the behavioral state could transition to one of two other states, so another set of parameters determines the state transition fractions (*σ*). The locations of resources are described by separate point sets, and movement among point sets for surviving mosquitoes during a time step is described by matrices (Ψ). Each adult module is made up of the state variables and specified by: point sets describing resources; parameters describing survival, success, state transitions; and movement matrices.

The models for adult mosquito population dynamics are coupled to a model for aquatic mosquito populations in the aquatic habitats. Success for gravid mosquitoes is egg laying, and a parameter describes the number of eggs laid (*o*). A model for aquatic mosquito population dynamics determines maturation, survival, and emergence. Emergent adults join the adult mosquito population ready to blood or sugar feed.

We developed a software package for R that takes advantage of the modular nature of mosquito ecology: adult and mosquito populations interact through terms that compute egg laying and emergence. The algorithms are encoded in an open source GitHub R package, called ramp.micro (Available at https://dd-harp.github.io/ramp.micro/). The code also includes algorithms to set up, solve the models, and analyze and visualize the outputs. The code to generate all the figures is available from https://github.com/dd-harp/dispersal_in_context.

Here, we present methods to analyze mosquito movement in two model families describing adult mosquitoes. In the first, called *BQ*, the adult population is either blood feeding (*B*) or egg laying (*Q*). In the second, called *BQS* (see Fig. 1), adults are either blood feeding (*B*), egg laying (*Q*), or sugar feeding (*S*). In the models, adult mosquitoes are located at a point where resources can be found.

**Fig 1.**
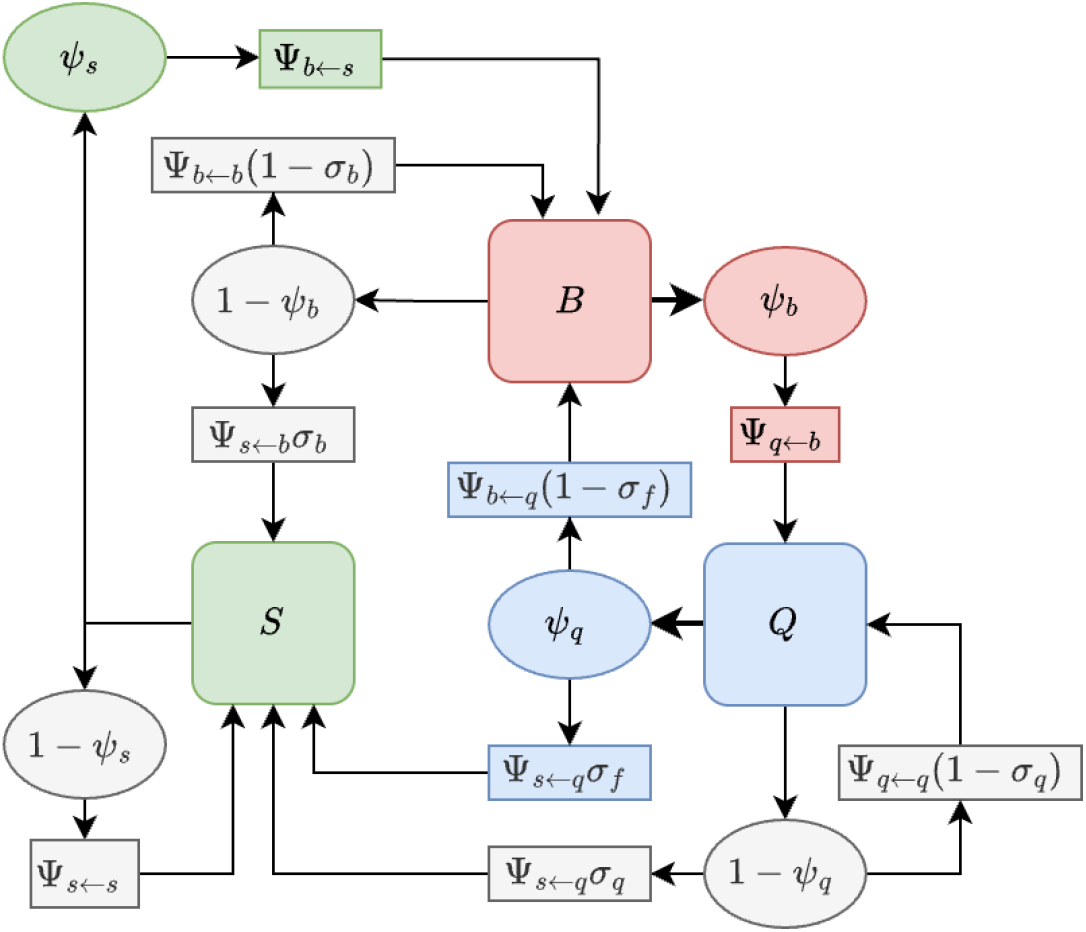
A diagram of the BQS model for adult behavioral states (squares) and state transitions (ovals). The rectangles describe transition probabilities. Adult mosquitoes are either sugar feeding (*S* located in 𝒮, green), blood feeding (*B* located in ℬ, pink), or egg laying (*Q* located in 𝒬, blue). After emerging, a proportion of mosquitoes will either blood feed or sugar feed (not shown). After a blood meal (an attempt succeeds with probability *ψ*_*b*_), a mosquito will always lay eggs. After successfully sugar feeding, all mosquitoes seek blood. After laying eggs, a fraction of mosquitoes sugar feed (*σ*_*f*_). After failing to lay eggs, a fraction of mosquitoes sugar feed (*σ*_*q*_), and after failing to blood feed, a fraction mosquitoes sugar feed (*σ*_*b*_). After changing states, a mosquito must also move to another point set, (from *x* to *y* is denoted by Ψ_*y*←*x*_).

In simulations, we found that the models are capable of a wide range of complex behaviors, depending on the resource landscape and parameters. In all of them, dispersal appeared to play a key role, so we developed methods to help us understand mosquito population movement. While each model describes movement in a single bout, we were interested in mosquito movement through multiple bouts to blood feed, sugar feed, or lay eggs; mosquito movement through one full feeding cycle; and lifetime mosquito movement. We also describe the spatial structure of mosquito movement. We describe the methods for analyzing movement here, and we invite users to explore the dynamics in greater detail using the software.

### Population Dynamics

We illustrate the analysis of movement using a *BQ* model with 256 blood feeding sites (denoted by *B*) and 289 egg laying sites (denoted by *Q*) that were randomly uniformly distributed. We also developed a *BQS* model with the same blood feeding and egg laying sites and 225 sugar feeding sites (denoted by *S*). Each one of the aquatic habitats was identical in every way, yet at the steady state, the population densities at the sites were heterogeneous (Fig. 2). Since there were no other differences, the heterogeneity was caused by dispersal patterns.

**Fig 2.**
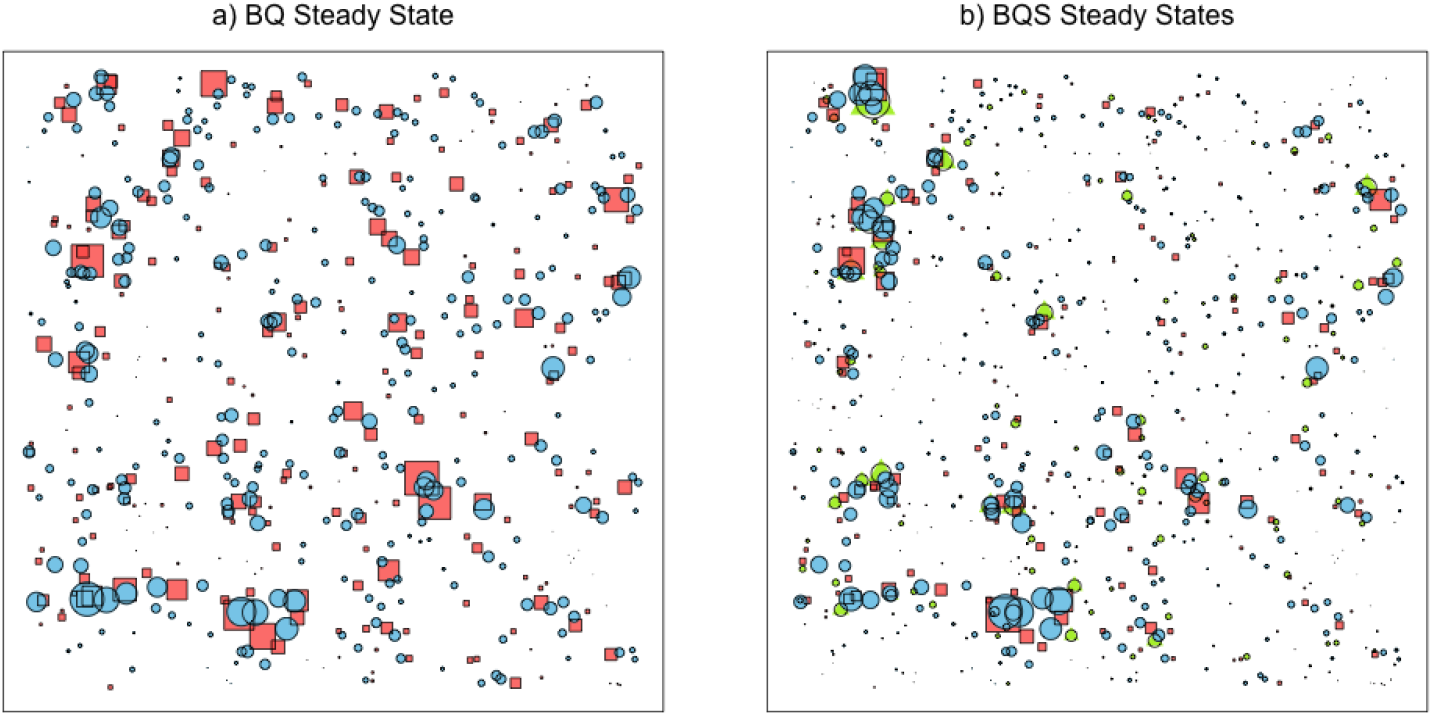
The population densities at the steady state for a) the *BQ* model and b) the *BQS* model, given the same resource distribution across the landscape. The red squares are blood feeding sites, ℬ; the size is scaled to steady state population densities. The blue circles are aquatic habitats 𝒬, scaled to steady state population densities. The green triangles are sugar sources 𝒮, scaled to steady state population densities. Aquatic dynamics are identical in every site, so all the variability in population density is due to the position in a network.

### Dispersal

In micro-simulation models, resources are located at points in space. Three resource point sets were defined: haunts where mosquitoes rest and where a mosquito might find and feed on a vertebrate host, a point set denoted ℬ, with *N*_*b*_ elements where *b*_*i*_ ∈ℬ describes the location of the *i*^*th*^ element; aquatic habitats where mosquitoes lay eggs, a point set denoted 𝒬 with *N*_*q*_ elements where *q*_*i*_ ∈𝒬describes the location of the *i*^*th*^ element; and possibly sugar sources where mosquitoes sugar feed a point set denoted 𝒮 with *N*_*s*_ elements where *s*_*i*_ ∈𝒮 denotes the location of the *i*^*th*^ element.

Movement among point sets is modeled using matrices that describe where mosquitoes move in a single flight bout, once each time step. The proportion moving from a point in one set to another, from each element in *x* ∈ 𝒳 to each element in *y* ∈ 𝒴, is described by a matrix Ψ_*y*←*x*_ or equivalently Ψ_*yx*_. Similarly, the proportion moving from a point in one set to another point in the same set is Ψ_*x←x*_ or Ψ_*xx*_ (the diagonal describes the probability of staying). The matrices describe the destinations for surviving mosquitoes, so each column sums to one. Mortality associated with dispersing is associated with the source points, and included in *p*_*x*_.

At the level of a flight bout, mosquito dispersal among point sets is defined by the matrix Ψ. For mosquito populations, dispersal to blood feed, lay eggs, or sugar feed can be understood through analysis of these population dynamic models, where a single task may require multiple attempts and incur multiple hazards. We have thus developed algorithms that compute the fraction surviving and dispersing through a single phase of the feeding cycle, or through one complete feeding cycle.

#### Movement to Complete a Task

Movement to accomplish a task could involve a sequence of search bouts, some of which may be unsuccessful. While we formulate the models with matrices describing movement in a single bout, a functionally relevant measure of dispersal tracks movement through one part of the feeding cycle (Fig 3): starting after laying eggs through some failures until a surviving mosquito successfully blood feeds, *K*_*b*←*q*_; or starting after blood feeding through failures until a mosquito successfully lays eggs *K*_*q*←*b*_.

**Fig 3.**
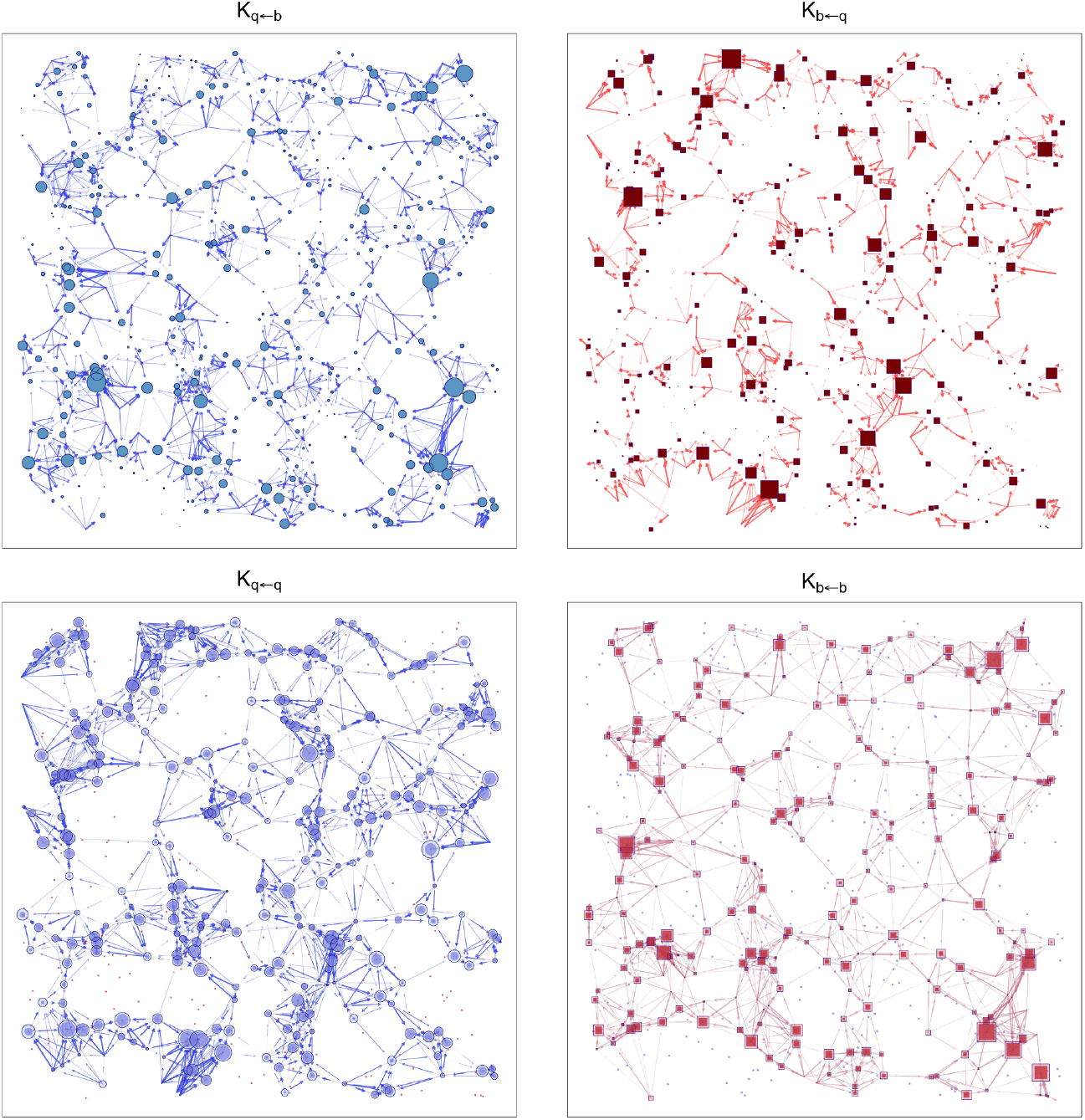
Movement to lay eggs or blood feed (top row) creates heterogeneity and structures movement among aquatic habitats or blood feeding sites (bottom row). **top row)** For one particular model, after defining the movement matrices, Ψ, and all the other parameters, we computed *K*_*b*←*q*_ (starting from points *q* ∈ 𝒬 and dispersing to points *b* ∈ ℬ to find blood); and *K*_*q*←*b*_ (*vice versa*, to lay eggs). **bottom row)** From *K*_*b*←*q*_ and *K*_*q*←*b*_, we computed matrices that show how mosquitoes move through one feeding cycle: *K*_*b*←*b*_ (from blood meal to blood meal among points in ℬ); and *K*_*q*←*q*_ (from egg batch to egg batch among points in 𝒬). In this visualization, the width of each *edge* is proportional to the propensity to move, and weaker connections are not plotted. The size of the points is proportional to mosquitoes arriving at each point. Asymmetric flows among pairs of points are represented using arrows.

Since there could be several failures, we modified the models to simulate one part of the feeding cycle. To do so, we modify the equations by setting parameters to zero that determine the fraction of mosquitoes that would otherwise leave the state. We then initialize and follow a cohort, iterating until almost surviving mosquitoes are trapped in the end state (*i*.*e*., up to a predefined tolerance). We let *K*_*b*←*q*_ denote a *N*_*b*_ × *N*_*q*_ matrix describing net dispersal to blood feed once: it is the proportion of mosquitoes leaving *q* ∈ 𝒬 after laying eggs that survive and eventually blood feed successfully at each point in *b* ∈ ℬ. Similarly, we let *K*_*q*←*b*_ denote a *N*_*q*_ × *N*_*b*_ matrix that describes net dispersal to lay eggs after blood feeding.

The matrices generated by the algorithm describe movement through blood feeding and egg laying, in models with or without sugar feeding. With sugar feeding, a mosquito that has laid eggs *must* blood feed before laying again. A mosquito may fail to lay eggs and switch to sugar feeding, and then blood feed again before successfully laying eggs (Fig 1). We can compute the matrices *K*_*b*←*q*_ and *K*_*q*←*b*_ accounting for the fact that a mosquito may sugar feed – possibly multiple times – before ending in the other state. Under the assumptions of this model, there may be some gonotrophic disassociation; a mosquito will always blood feed after sugar feeding, even if it was searching for an aquatic habitat before the transition (Fig. 1). In such cases, *K*_*b*←*b*_ is the interval between blood meals with or without an egg batch, but because of the delay for parasite development, we must take even greater care in quantifying parasite dispersal by mosquitoes (see below).

#### Movement through a Feeding Cycle

After computing dispersal through one part of the feeding cycle, the matrices *K*_*bq*_ and *K*_*qb*_ define a new, bipartite graph that describes movement among the point sets where mosquitoes lay eggs and where they blood feed (see Fig 3). Movement through egg-laying and (possibly) sugar feeding can be represented in block form:

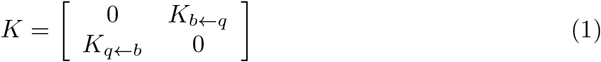

After blood feeding and egg laying once – one complete feeding cycle – the bipartite graph disconnects into two complementary graphs on their respective point sets:

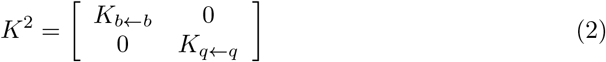

The matrix *K*_*b*←*b*_ = *K*_*b*←*q*_ · *K*_*q*←*b*_ describes movement from haunts to haunts, and *K*_*q*←*q*_ = *K*_*q*←*b*_ · *K*_*b*←*q*_ describes movement from habitats to habitats (Fig. 3).

#### Egg Laying

Next, we developed algorithms to compute net dispersal of eggs away from a natal site (see Fig 4). Net lifetime dispersal of eggs from natal aquatic habitats is summed over all gonotrophic cycles, *i*, until a mosquito has died. If mosquitoes lay *o* eggs per batch, the expected number of eggs laid per emerging adult female at each site can be computed as:

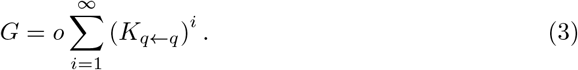

**Fig 4.**
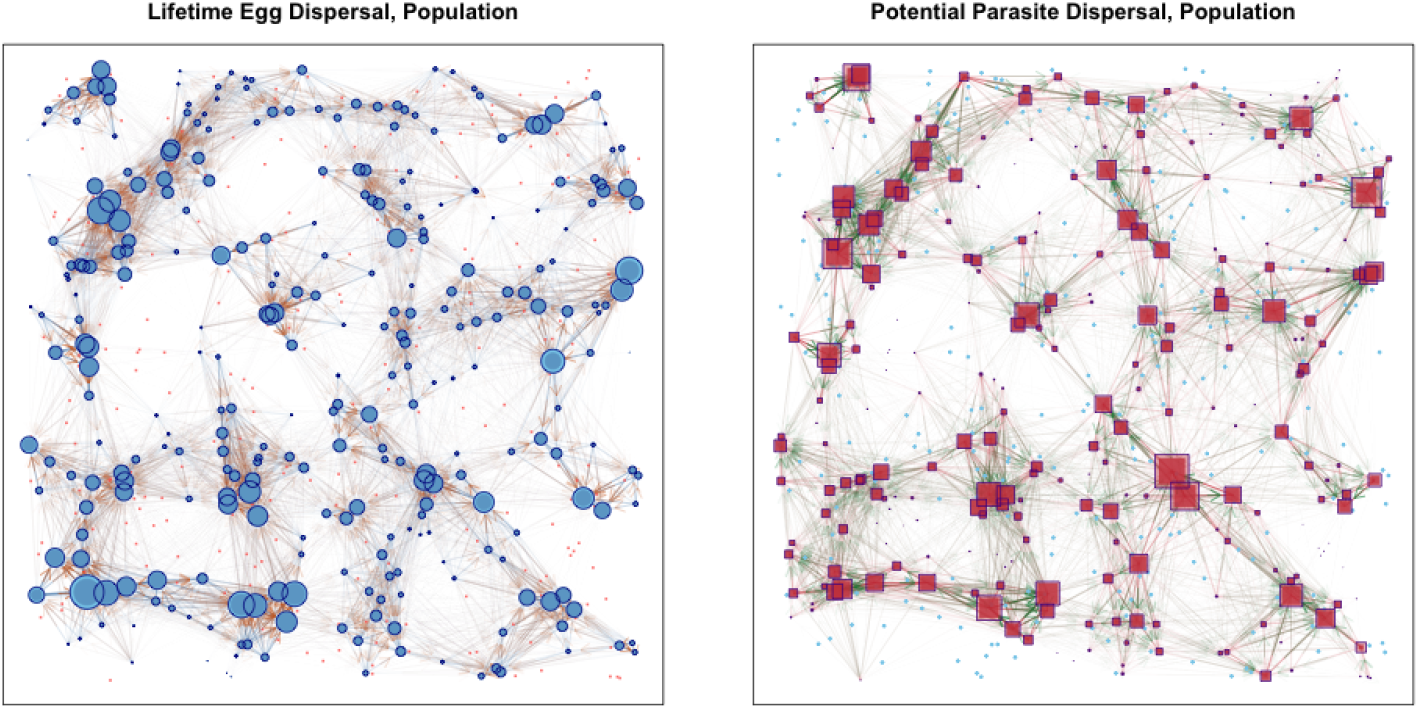
For the model in Fig 2a and 3, these plots show lifetime egg dispersal by mosquito populations (𝒢), and potential parasite dispersal by mosquito populations (𝒱). The size of each point *q* ∈𝒬 (dark blue, top) and *b* ∈ ℬ (dark red, bottom) is proportional to the number of infectious bites or eggs laid *arriving* at each point (*i*.*e*. not arising from). A key feature of these graphs is that some haunts or habitats tend to *receive* more infective bites or more eggs because of topology, *i*.*e*. their position in the graph describing movement. In this visualization, the width of each *edge* is proportional to the propensity to move, and weak connections are not plotted.

The quantity *G* is mosquito lifetime egg dispersal kernel for a cohort. We also define net egg dispersal by the population at its steady state

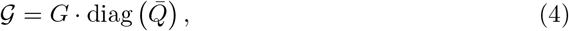

where 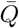 is the steady state density of adult female mosquitoes at *q* ∈ 𝒬.

#### Pathogen Dispersal

We developed algorithms to compute potential dispersal of pathogens from the points where mosquitoes blood feed (Fig 4). A basic formula for the dispersion of infective bites away from an infectious host must consider a delay for the pathogen development and a latent period, traditionally called the extrinsic incubation period (EIP). Since chronological and reproductive age are not necessarily in sync, and since there might be gonotrophic disassociation, we can’t use *K*_*b*←*b*_. Instead, we simply run the simulation, summing up all potential transmission events, just as we would compute vectorial capacity.

Net potential dispersal of pathogens is computed by setting an initial cohort of mosquitoes that successfully took a blood meal, *B*_0_, among haunts and iterating the model until all of the mosquitoes are dead. The number of infective bites given by the cohort is found by adding up all the bites after *τ* days, where *τ* is a parameter describing the EIP. Given *B*_0_, we use the model to compute *B*_*t*_ for a cohort of mosquitoes, and we define *V* as

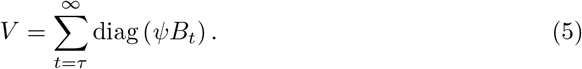

While *V* defines parasite dispersal by a cohort of mosquitoes among haunts, where each point counts equally, we can also weight parasite dispersal by the steady state population density at each point, which we call potential pathogen dispersion,

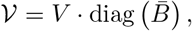

where 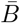 is the steady state density of adult female mosquitoes at *b* ∈ ℬ.

The matrix 𝒱 defines a quantity that differs slightly from the definition of vectorial capacity, in that it is the expected number of bites arising from all the mosquitoes blood feeding at a haunt, rather than on a single person, computed as if every available host was perfectly infectious. The quantity would be identical to *vectorial capacity* if it were normalized by the number of hosts available (*i*.*e*. modifying *B*_0_), such that the quantity was an expected number arising from a single host. Note also that *V* and 𝒱 an be used to computes bites arising from or arriving at each haunt by summing over rows or columns.

#### Dispersal Distances

Any pair of matrices describing dispersal propensities among points in a set and the distances among points in that set can be normalized to define a probability distribution function. Since the number of destinations is finite, dispersal is described using probability mass functions (PMFs).

### Population Structure

The matrices describing mosquito movement among point sets can be understood through the study of the associated graphs: resources are represented by vertices, and movement propensities are represented as directed, weighted edges with a self-loop. We investigated *community structure* by converting dispersal matrices into graphs and analyzing the results using methods developed for networks. Since most algorithms are designed to analyze simple graphs, we used the walktrap algorithm from *iGraph*, which can do network analysis for directed, weighted graphs. (Other community detection algorithms could be used, but since walktrap analyzes clustering based on short random walks, it seems to be a good fit for analyzing flights by short-lived mosquitoes.) The algorithm returns a set of communities, groups of points that are more densely connected to each other than to the rest of the points. Information about location is used to develop the dispersal matrices, and location is also used to plot and visualize the results, including the communities identified from analysis of the graphs.

Initially, we sought to analyze the community structure for crude dispersal matrices (for the *BQ* model, see Eq. 13). The comparison with our other analyses were often at odds. The main difference is that since the mosquito populations are structured by the feeding cycle, a sequence of mosquito flight bouts is not switching randomly among states, but structured by the feeding cycle, from one task to another (Eq. 1). Movement through the feeding cycle is structured by bipartite graphs, and after a feeding cycle, the bipartite graph separated into two independent graphs (Eq. 2) – one for egg laying among aquatic habitats, and another for blood feeding among haunts. Community analysis for crude dispersal matrices (*e*.*g*., Eq. 13) did not agree with the analysis for the *K*^2^ matrices (Eq.2), in part, because mosquito biology imposes some order to the kind of random walks a mosquito takes.

This discrepancy suggests that a crude analysis of observed mosquito movement patterns might not reveal community structure unless it could somehow account for mosquito behavioral states.

#### Communities

Our first point sets were generated using randomly uniformly distributed parameters, yet the resulting graphs for describing movement for these basic point sets displayed a high level of organization (see Fig. 4). The co-distributions of most pairs of point sets will have some degree of clustering. In these models, the patterns get amplified by the movement rules. Mosquito populations were self-organized into communities around places that had all the required resources nearby. Not surprisingly, the community structures for eggs laid and parasites dispersed tended to overlap geographically (Fig. 5). By studying the flows on these bipartite graphs, we began to understand what generated patterns in the steady state densities of mosquitoes at haunts and habitats.

**Fig 5.**
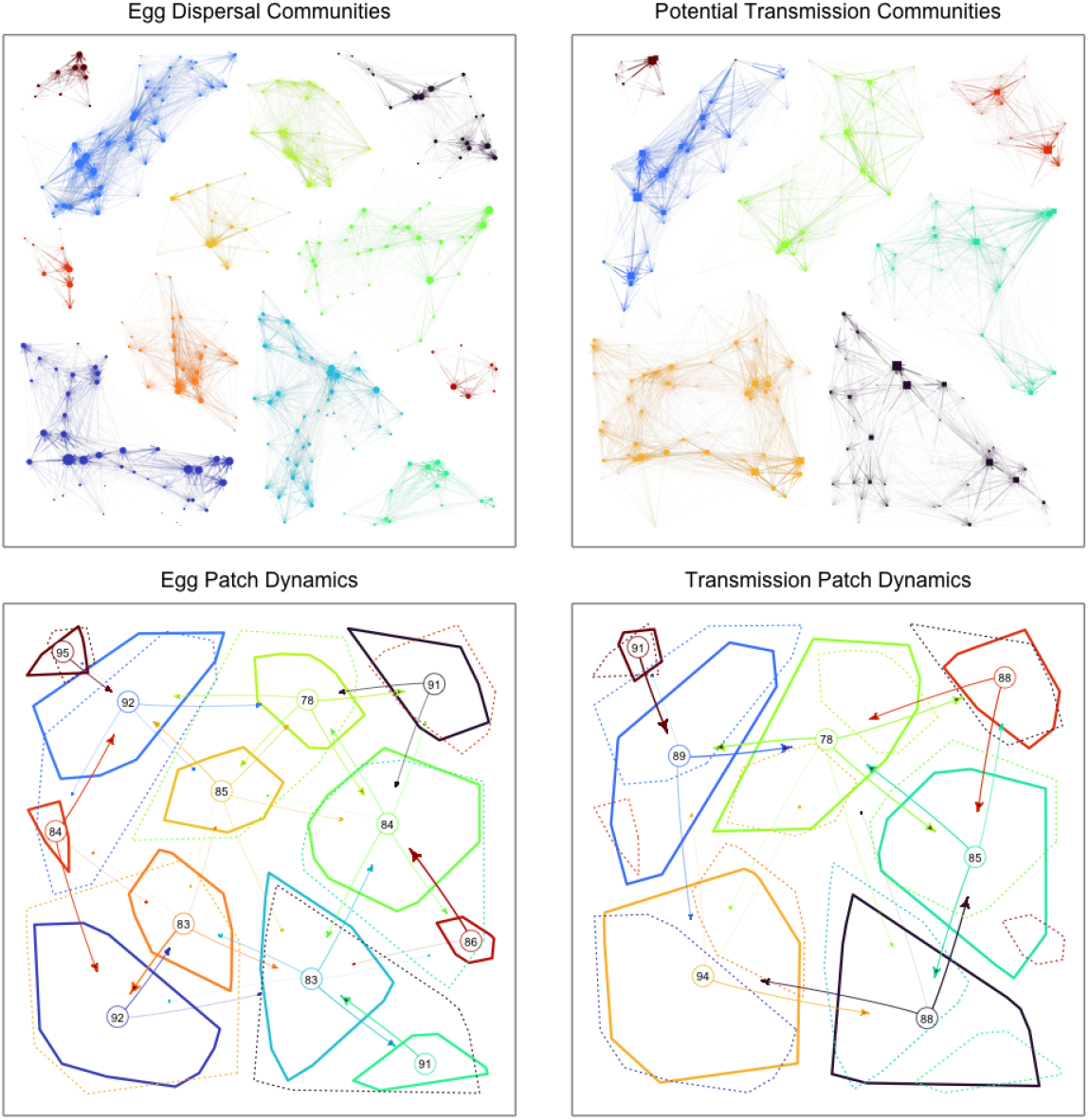
For the dispersal graphs in Fig. 4, we have plotted a subgraph for each community that was identified by the walktrap algorithm (from *iGraph*) for egg laying (left, 𝒢) and potential transmission (right, 𝒱). Each community sub-graph is given a different color After aggregating flows within and among communities, the arrows show the net flows within each *patch*, plotted as a convex hull around the community. The convex hulls from the complementary process are also plotted in a lighter color in dashed lines. The number at the center is the percentage of the total that remains in each patch. The bent edges show the proportion dispersing out from each patch to every other patch.

These communities could serve as spatial *patches* in a patch-based meta-population model. Even though we started from a different mathematical formalism, the assumption commonly made for patch-based models – that a constant fraction of mosquitoes in one patch move to other patches – would approximate the processes reasonably well. Notably, any set of patches would approximate the underlying process to some degree: there would be more movement occurring within *vs*. among communities would also work for *any* tessellation of the plane. The critical difference is that the meta-population dynamics would approximate the underlying processes better if the patches reflected community structure.

#### Flows

In analyzing dispersal through a feeding cycle, we noted that the heterogeneous distribution of resources created a propensity for mosquito populations to *flow* among point sets. The mechanism generating the flow was related to asymmetries in the *relative* position of points; after leaving a resource at one point in space, finding another resource, and launching a new search from there, the propensity to return to a focal point depended on the arrangement of all the other points nearby. The use of a resource of one type depended on its relative proximity to resources of the other types. For example, in the *BQ* model, the points *b* ∈ *B* that are most likely to be found starting from *q* ∈ 𝒬 are not always the same as the points in *q* ∈𝒬 that are most likely to be found from each starting point *b* ∈ ℬ. Given the opportunistic nature of searching, mosquitoes would tend to stay in areas where all the required resources can be found, and mosquitoes would tend to leave the neighborhood of resources in one set that are not comparatively close to other resources in the other. These asymmetries in movement propensities can be visualized in the graphs of *K*_*b*←*b*_ and *K*_*q*←*q*_ (Fig. 3). Even in point sets generated using uniformly distributed random numbers, these relationships are rarely symmetrical. The arrangement of points robustly introduces asymmetries that create a propensity for mosquito movement graphs to flow away from some areas and into others.

In analyzing the community structure (see below), we had concerns about the nature of the asymmetric flows and the validity of the analysis of community structure. It is possible, for example, that the net flows in some of these graphs would be so strong and important that they could undermine the cohesiveness of any communities that could form. A graph of turbulent flows of water in a river, for example, could identify important eddies that reflected a kind of transient structure that was constantly broken down by a downstream flow.

After trying several different ways of describing or quantifying patterns, we found the greatest insights from comparing the mosquitoes arriving at each point after a single feeding cycle (*i*.*e*., from *K*_*q*←*q*_ and *K*_*b*←*b*_) to population densities at the steady state. The two distributions were similar, which was not surprising, because in these models, population density at the steady state was higher *because* more mosquitoes moved there. The patterns that emerge in populations reinforce the patterns generated during a feeding cycle. Haunts or habitats with low population densities were found at the periphery of a community, while those with high densities formed the core. Perhaps more relevantly, we did *not* find any features of the flows – something like the directional flow resembling a river – that would give rise to concerns about the soundness of the network analysis. Instead, we found that the asymmetric flows appear to create and reinforce community structure.

### Context and Pattern

In developing methods for analyzing mosquito movement on complex resource landscapes, we chose a single landscape where the resources were randomly, uniformly distributed. The analysis of movement in Figs 2-5 all apply to this landscape. The kind of structure that emerged is not a feature of most diffusion-based approaches to understanding mosquito dispersal (but see [13]). This natural patchiness could, perhaps, be handled just as well in metapopulation models where mosquitoes emigrate from a patch at a higher rates when a required resource is unavailable [1]. The framework points out the need for more work to understand the structural limitations of the models we use to describe mosquito ecology as complex systems, but that is beyond the scope of this study.

These behavioral state microsimulation equations are one framework for developing realistic agent-based models of mosquito populations that are easier to analyze than individual-based simulation models [15]. In this framework, mosquito dispersal has a purpose: mosquitoes fly around searching for resources. The movement patterns of mosquito populations are thus structured by those resources. The patterns emerge from searching for different resources in a sequence that follows a mosquitoes behavioral programming. The framework is realistic enough that it might be possible to rigorously define the concept of a mosquito’s fundamental niche as an emergent feature of 1) a mosquito’s behavioral algorithms; 2) the quality, stability, and suitability of aquatic habitats; 3) the availability of other resources; and 4) the costs of searching.

Any framework that is capable of defining a mosquito’s fundamental niche would also describe resource landscapes where mosquitoes would not persist. With enough imagination, patterns of all sorts could emerge. Any competent mosquito ecologist could use these models to understand how context shapes mosquito population dynamics, but the approach would require development of new approaches to studying mosquito populations, perhaps one that is more strongly grounded in evolutionary ecology.

While mosquito population structure can form even if the underlying distribution of resources were drawn from a uniform random distribution, the claim is that mosquito population structure emerges because searching mosquitoes to leave an area while searching for a resource that is not available. To test this, we simulated mosquito populations on a modified version of our landscape by restricting the distribution of one or more resource.

If the population persists, mosquitoes will tend to be aggregated around resources that are rare. To illustrate, we created new landscapes, derived from the original, where one of the resources was rare (Fig. 6). These correspond roughly to a rural focus, where aquatic habitats are common but humans are rare (Fig. 6a); an urban focus, where humans are abundant, but aquatic habitats are rare (Fig. 6b); and finally, a model that demonstrates how the distribution of sugar could explain adult mosquito population distributions, even when human blood hosts and aquatic habitats are common.

**Fig 6.**
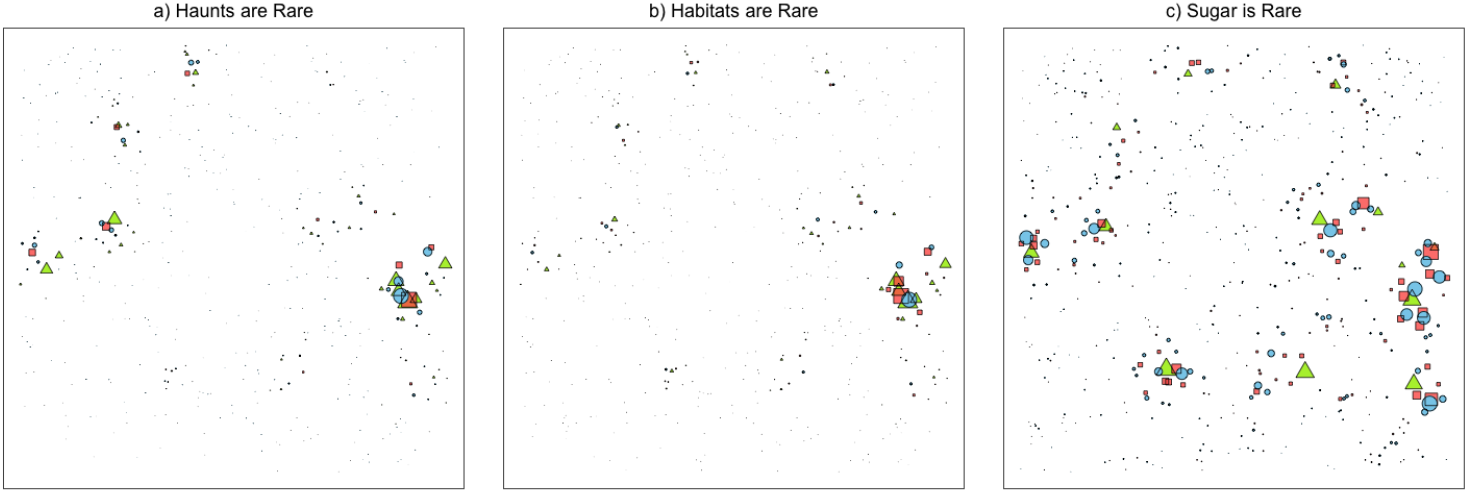
A set of simulations in which each point set in Fig 2b is replaced by a point set with only 13 points. The effects on population dynamics are different in each case. Three of these points are reasonably close to each other, and the proximity seems to amplify the local population dynamics. Note that mosquito populations cluster around the point set with least common resource. Rare habitats have the strongest effects on mosquito population structure. Since sugar is an optional resource – mosquitoes can blood feed opportunistically – the effect of sugar being rare is not as strongly limiting.

## Discussion

Using models that mimic some basic mosquito behaviors, we have described how mosquito movement is shaped by context, i.e., the spatial distribution of resources on a landscape. In these models, mosquito population movement patterns emerge from simple behavior rules and the joint distributions of resources. This presents a challenge for the study of mosquito dispersal, because if context matters, then what general lessons can be learned from developing and analyzing movement on any particular landscape? On the other, context dependency is probably a feature of real systems, which are challenging and expensive to study [2], so there is a great advantage to learning about these systems through the study of *in silico* systems.

Population structure in these models emerges from the co-distribution of essential resources and mosquito searching behavior, and it gives rise to heterogeneity and patchiness in the distribution of adult mosquito populations. A surprising amount of fine-grained population heterogeneity and structure arises from applying simple rules to simulate search and dispersal on simple heterogeneous resource landscapes. The patches are self-organized around locations that have all the essential resources that mosquitoes require to survive and reproduce. Conversely, places on the landscape that lack one or more of a mosquito’s essential resources will tend to have lower densities as mosquitoes leave while searching. The likelihood of returning to a spot would depend on the arrangement of the other resources. A resource will get used if it is close to other required resources. After leaving one resource in search of others, and reverting to searching for that resource again, the mosquito may be more likely to find it at another location that is closer. The resulting probabilistic process creates a tendency for mosquitoes to move away from areas that lack some resources and stay in areas that have all of them. Mosquito populations will thus tend to be aggregated in places that have all the resources a mosquito requires.

Mosquito communities, defined by mosquito connectivity in networks describing mosquito movement among resources, are a self-organized spatial phenomena that resembles the patches in meta-population models. Even within these patches, where all resources are available, the co-distribution of resources and mosquito search tends to produce *flows* among resources that are a cause of heterogeneity. There is substantial mosquito movement within a community as mosquitoes fly back and forth among resources, with weaker connectivity among patches as mosquitoes disperse long-distances among communities. Because of wind and other factors that give shape to the function that we used to compute dispersal matrices, the length of flight bouts could be Levy-like if we used fat-tailed functions, so that mosquito populations would be weakly coupled across fairly long distances. A core difference is that with metapopulation models, the patches are defined arbitrarily, not as an emergent feature.

The study of behavioral-state microsimulation models for mosquito ecology is challenging. Given the importance of contextual factors and the difficulty of systematically exploring the dynamics, we have demonstrated the importance of context, but we would hesitate to make any general claims about mosquito community structure. To facilitate the study of these models, we developed an R package. This software can lower the costs of developing and analyzing such models, and makes these studies more replicable. The software is extensible, making it possible to add features in the future to address some of the limitations of models we have developed: aquatic habitat dynamics can change the structure of mosquito populations seasonally; search involves wind, so the speed and direction of the wind probably shapes mosquito dispersal; topography affects how mosquitoes caught by the wind end up landing in specific areas, in the lee; in real systems, mosquito search is probably affected by specific features of a landscape; mating is an important feature of mosquito population dynamics. These are potential areas for software development and future studies.

Perhaps most importantly, these methods are realistic enough that we can begin to develop a quantitative basis for defining a mosquito *niche*. A mosquito population can only persist in places that have all the essential resources located close enough to each other that a mosquito can complete its feeding cycle and lay eggs well enough for mosquito populations to persist. We suspect an important feature of real systems is a mortality risk associated with searching [28], but it could be useful to explore how migration from an unfavorable starting location or under unfavorable conditions could lead to mosquito mortality, raising other questions: How much does a mosquito use environmental cues, such as relative humidity, wind, or sun to determine when to launch a flight bout? What search algorithms would optimize mosquito fitness on particular resource landscapes? Mosquito population dynamics are an emergent feature of the organism’s needs, behavioral states, behavioral algorithms, and the environment as it minimizes hazards and searches for resources distributed heterogeneously on landscapes. These models could be modified to consider how mosquitoes respond to environmental cues and how they are affected by specific hazards. Environmental factors such as humidity, temperature, and wind could affect when a mosquito decides to launch. The availability of resources structures how they disperse. When these are combined with models describing aquatic dynamics, there is a sound basis for a grand unified theory linking mosquito genetics and physiology to mosquito population ecology in some context. The approach is amenable to evolutionary ecological analysis to understand how the behavioral algorithms of mosquito populations could adapt to local conditions. All of his suggests that behavioral state micro-simulation could provide a framework for a new synthesis.

Despite the formalism and the many ways mosquito behavior is not like diffusion, our analysis suggests that patchiness emerges, so patch-based meta-population models could capture much of what is important about that spatial structure. Patch-based models could be good approximating models for mosquito movement in some cases. Using the concept of resource availability to understand heterogeneous emigration rates by state, metapopulation models could generate the same sorts of heterogeneous mosquito dispersal and flows among patches [1]. The patterns that arise in these highly mimetic models raise questions about whether other emergent features could arise when considering stochastic dynamics or realistic aquatic dynamics. The study calls for a re-examination of the validity of the formulas used to estimate mosquito population size using mark-release-recapture data, as fine-grained structure could introduce substantial bias [29, 30]. Further ecological investigation is also warranted to explore the ability of mark-release-recapture studies with multiple release points [31], or close-kin mark-recapture studies [32], to detect emergent mosquito patches within real landscapes with naturally heterogeneous distributions of resources.

These models that start from *realistic* assumptions about resources and search provide an alternative way of thinking about and studying mosquito movement and its relevance for other problems in mosquito ecology and mosquito-borne pathogen transmission. The predictions of these micro-simulation models could be used to study the spread of genetically modified mosquitoes with frequency dependence [33], and they hint at ways that mosquito movement could affect micro-dynamics of mosquito-borne pathogen transmission. Similar approaches can inform the placement of traps for the design of mosquito surveillance systems [34]. While we used these models to study potential transmission, they have not (so far) included any vertebrate host infection dynamics. Notably, theory for pathogen transmission by mosquitos suggests that potential transmission intensity is proportional to the ratio of mosquitoes to humans; it is directly proportional to mosquito density and inversely proportional to human population density, but there has been very little consideration of the relevant spatial scales for computing the ratios. One hypothesis suggested by these models is that the propensity for mosquitoes to cluster around places where resources are available creates something like a *focus*, an area where transmission could be sustained in a small area involving a small number of hosts.

Despite the context dependency and complexity, the lessons learned could prove to be generally useful. Importantly, the use of one resource tended to depend on its proximity to the other required resources. Our analysis suggests that mosquito populations and potential transmission will tend to cluster around places where all required resources are available. Heterogeneous resource distributions – in particular the relative paucity of one or more required resources – give rise to complex, structured, heterogeneous mosquito populations. The hope is that at least some of these ideas could give rise to methods for more effectively targeting interventions to reduce the burden of diseases and control transmission.

## Acknowledgments

Funding for this project for HMSC, SLW, JMY, and DLS was provided by grants from the Bill and Melinda Gates Foundation, OPP 1159934, OPP 1110495, and INV 030600. Funding for DLS was also provided by an grant R01AI16339 from the US National Institutes of Allergies and Infectious Diseases. JMM was supported by a National Institutes of Health R01 Grant (1R01AI143698).

## Methods

The software to set up models, run simulations, analyze patterns, and visualize ouptuts for these systems is available as an R package on GitHub, called ramp.micro (https://dd-harp.github.io/ramp.micro/). The software includes utilities to build resource landscapes, to compute various summary statistics describing mosquito population dynamics and mosquito and pathogen dispersal, and to visualize networks and communities.

### Behavioral State Dynamics on Point Sets

We define micro-simulation models on point sets representing the locations of resources: haunts where mosquitoes rest and where a mosquito might find and feed on a vertebrate host ; aquatic habitats where mosquitoes lay eggs (a point set denoted *Q* with elements *q*_*i*_ where *i* = 1, 2, …, *N*_*q*_); and possibly sugar sources where mosquitoes sugar feed (a point set denoted *S* with elements *s*_*i*_ where *i* = 1, 2, …, *N*_*s*_). In the simulation models, the number of adult mosquitoes emerging from each aquatic habitat on each day is a vector denoted Λ_*t*_ that can be either passed as a parameter or simulated using a population dynamic model for aquatic immature population dynamics (see below). Adult mosquitoes are located at one of these points at each point in time (*e*.*g*., one day), represented as a set of vectors: *B*_*t*_, the number seeking blood at haunts; *Q*_*t*_, the number attempting to lay eggs at habitats; or *S*_*t*_, attempting to feed at sugar sources. Mosquito bionomics can depend on both behavioral state and location, including daily survival (vectors denoted *p*_*b*_ or *p*_*q*_ or *p*_*s*_), and daily blood feeding or egg laying success (*ψ*_*b*_ or *ψ*_*q*_ or *ψ*_*s*_). For convenience in writing the equations, below, we use a symbol to denote the complement: 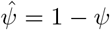.

Movement among point sets is modeled using matrices that describe where mosquitoes move each time step. The proportion moving from a point in one set to a another, from *x* ∈𝒳to *y* ∈ 𝒴, is described by a matrix Ψ_*y*←*x*_ or equivalently Ψ_*yx*_. Similarly, the proportion of mosquitoes that are re-attempting a task that move from a point in one set to a point in the other is Ψ_*xx*_. The matrices describe where surviving mosquitoes end up; all mortality is described in the survival parameters (*p*). Since there are a finite number of destinations, each column is a probability mass function (PMF).

#### Immature Population Dynamics

For purposes of integrating the effects of movement and aquatic ecology, we assume each female lays *o* eggs, so the total number of eggs laid each day in each site is:

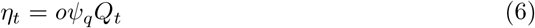

We assume the number of immature mosquitoes in aquatic habitats, denoted *L*_*t*_, is subject to density dependence, which could delay maturation or increase mortality. The maturating fraction is, 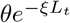 where *ξ* > 0, and surviving fraction is 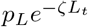 where *ζ* > 0. The parameters are site-specific, so that some habitats can vary in quality. The dynamics are:

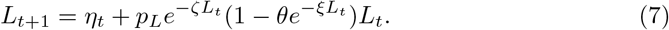

The number of adult females emerging each day is half the mosquitoes who both survived and matured:

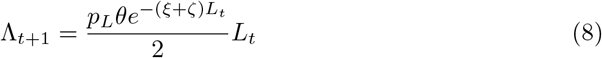

While this model is adequate for the needs of this study, the model may not robustly capture some relevant features of mosquito ecology or larval source management, such as when larval population structure and delays are important. The software, `ramp.micro`, includes alternative models with stage structure.

#### A Basic Feeding Cycle Model

In the basic feeding cycle model over one time step, mosquitoes either attempt to blood feed or attempt to lay eggs. The result of an attempt is either survival or death, and if the mosquito survives, success or failure. A success moves a mosquito to the other state, and if they fail, they must try again. Either way, the mosquito moves to a point in the set for the resource they seek: the diagonal of Ψ_*x*←*x*_ is the probability of staying. Using⟨·⟩instead of diag (.) for the diagonal matrix, the equations are:

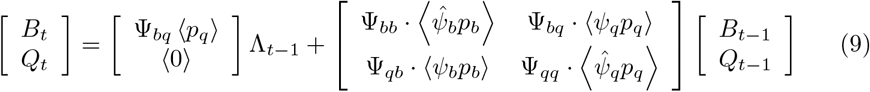

where Λ_*t*_ is the number of emergent adult female mosquitoes on day *t* (see above).

#### A Feeding Cycle Model with Sugar Feeding

In the models with sugar feeding, the updating basic rules are similar, but a new set of parameters are required to describe the frequency of switching to sugar feeding from other states. Each day, some fraction of mosquitoes switch to a sugar feeding state from various points in the feeding cycle (Figure 1): the switch to sugar feeding occurs in a fraction of recently emerged mosquitoes *σ*_Λ_; after egg laying *σ*_*f*_ ; after a failed egg laying attempt, *σ*_*q*_; or after a failed flood feeding attempt, *σ*_*b*_. We assume that all mosquitoes revert to blood feeding after sugar feeding, implying that the mosquito resorbed the eggs. We also assume that mosquitoes would never attempt to sugar feed after a blood meal, but that they would instead always attempt to lay eggs at least once (Fig. 1).

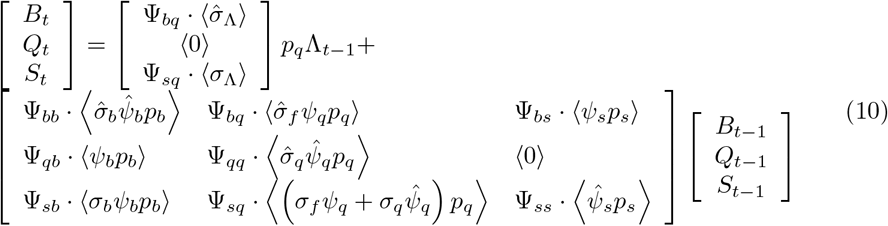

### Resource Landscapes and Dispersal Matrices

To simulate mosquito dynamics, it is necessary to fully specify all its elements, including the parameters and resource point sets. Dynamics and patterns emerge from the co-distribution of those points and from the population processes. Aggregate movement patterns of mosquito populations in models of this type are determined by how many sites exist and which sites a mosquito ends up finding. Patterns emerge as movement and survival are iterated over multiple search bouts. To develop some analytical methods and some basic intuition, we start with randomly generated sets of points and simple distance-based movement rules.

We generated sets of points within a spatial domain using various probability distribution functions, including a uniform random distribution, lattices, and clustered points generated by various algorithms.

We use functions to generate the probability of finding a point as a function of its distance and context. Let *d*_*i,j*_ denote the distance from point *i* to *j*. We define some function to assign a weight to every pair of points. In this model, we use the exponential family of functions to assign search weights:

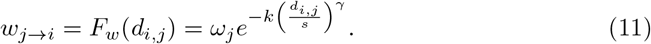

The scalar *ω*_*j*_ is a linear weight on each destination making it more or less attractive from every distance. In **ramp.micro**, other functional forms have been developed, including shapes that have a *fat tail*.

The search weights are normalized and translated into proportions: the proportion arriving at each point in a destination set index by *j* from a source set indexed by *i* is normalized across all destinations from each starting point:

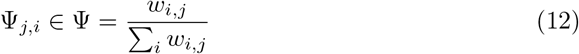

To be clear about the assumptions of this model, a matrix, Ψ, describes the probability of arriving at a destination conditioned on the mosquito surviving the flight, so the columns of Ψ sum up to one. Mortality associated with dispersal away from a point is included in the parameter describing daily survival of a mosquito *leaving from* each point. Each column of each Ψ thus defines a PMF for surviving mosquitoes, but statistics describing movement of mosquitoes depend on how many mosquitoes in the population start from each point and the resource they seek.

In the models we examine herein, the dispersal matrix is not time-dependent, but time-dependent matrices could be adopted to simulate other dispersal rules and effects, such as wind with variable speed and direction.

### Simulation

For any landscape, we can simulate mosquito population dynamics over time. If all parameters are constant over time (*i*.*e*. values of *p, ψ*, Ψ, *ξ, θ*, and *ζ*), then we can find a steady state either by inverting a matrix or by running the simulation for a long time. We let 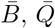, and 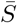 denote the steady state densities of adult mosquitoes in the point sets.

### Community Detection

We analyzed all of the square community matrices defined on single point sets, including *K*_*bb*_, *K*_*qq*_, *G, V*, 𝒢, and 𝒱.

We were also interested in analyzing the matrix describing the bulk movement of adult, female mosquitoes among all patches. For the model without sugar feeding, movement propensities by mosquitoes through one flight bout are described by the matrix:

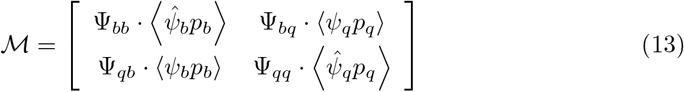

We also analyzed the closely related matrix that describes the movement propensities of mosquito populations, which is weighted by steady state densities:

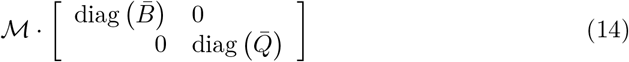

The matrix for the graph with sugar feeding is to Eq. 10 as Eq. 14 is to Eq. 9.

For any movement matrix, we examined population structure by analyzing these matrices as graphs. To summarize movement patterns, we identified “communities” defined by these graphs. A community expects more transitions (mosquito jumps) amongst the points within the community than transitions to a different community. Since the graphs are potentially asymmetric, we used the *walktrap* algorithm from the R package *iGraph*.

Notably, mosquito survival and dispersal work like the walktrap algorithm for community detection. The short lifespan of mosquitoes means that most mosquitoes will tend to move approximately 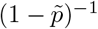 times in random walks, where 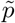 is something like a geometric mean lifetime probability of survival.

The graphs we analyzed were directed, weighted graphs with a self-loop. To understand these graphs, we decomposed them into two related graphs: the matrix *D* describing a graph can be expressed as the sum of a symmetric matrix *D*_*s*_ and an asymmetric matrix that represents the residual flows *D*_*f*_ . The symmetric matrix is the minimum of the matrix and its transpose, *D*^*t*^:

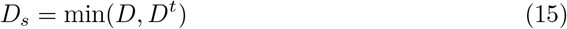

and the residual flow is:

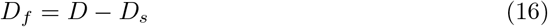

Note that for all *d*_*i,i*_ ∈ *D*_*f*_ that *d*_*i,i*_ = 0. If *i* ≠*d*_*ij*_, then for all pairs *i, j*, either *d*_*i,j*_ = 0 or *d*_*j,i*_ = 0.

We examined the residual flows created by *D*_*f*_ and *D*_*s*_ separately to better understand the nature of the mosquito population movement patterns that emerge from these simple rules.

To examine the robustness of the community structure, we run walktrap both on the graphs and on their decomposed matrices (*i*.*e*. for each graph *D* that we examined, we also examined *D*_*s*_ and *D*_*f*_). For an extended discussion, see the articles at https://dd-harp.github.io/ramp.micro/.

